# Genomic Landscape of Macrolide Resistance in High-Burden Bacterial Pathogens: Implications for Antimicrobial Stewardship in Resource-Limited Settings

**DOI:** 10.1101/2025.03.11.642729

**Authors:** Selassie Louis Ameke, Selinam L. Ameke, Sedem Agbovi

## Abstract

**Background:** Antimicrobial resistance (AMR) threatens global health, with macrolide resistance emerging as a critical challenge in high-burden pathogens. Understanding taxon-specific resistance mechanisms is essential for guiding interventions in resource-limited settings where diagnostic and therapeutic resources are constrained.

**Methods:** We analyzed 9,706 bacterial genomes from the AMRFinderPlus database to characterize macrolide resistance alleles, their distribution across taxa, and associations with multidrug resistance. Statistical and visualization workflows identified taxon-specific patterns and clinical implications.

**Results:** Macrolide resistance mechanisms dominated (34% of 1,243 alleles), with 23S rRNA mutations prevalent in *Neisseria gonorrhoeae* (23S_A2045G: 62%; 23S_A2057G: 58%) and *Streptococcus pneumoniae* (23S_A2058G: 48%). Efflux pump variants (acrB_R717L/Q) drove multidrug resistance in *Escherichia spp*. (41%) and *Salmonella spp*. (33%), correlating with cross-resistance to fluoroquinolones (18%) and tetracyclines (14%). These pathogens disproportionately impact regions like Ghana, where syndromic management and antibiotic misuse amplify resistance.

**Conclusions:** These findings underscore the urgent need for precision surveillance and context-specific stewardship to combat macrolide resistance in high-burden pathogens, particularly in resource-limited settings where diagnostic and therapeutic gaps exacerbate AMR threats. Targeted interventions, including affordable molecular diagnostics and antibiotic access reforms, are critical to mitigating global health disparities driven by resistance.

## Introduction

Antimicrobial resistance (AMR) is a global health crisis, but its burden is disproportionately borne by resource-limited regions, such as sub-Saharan Africa, where constrained healthcare infrastructure and diagnostic gaps exacerbate treatment failures and mortality (Otieno et al., 2022; Tornberg-Belanger et al., 2019). Macrolides, including azithromycin and erythromycin, are widely used in these settings due to their affordability and broad-spectrum activity against respiratory, gastrointestinal, and sexually transmitted infections (Almehizia et al., 2023; NUDANU and FAYED, 2023). However, the escalating prevalence of macrolide resistance driven by 23S rRNA mutations (Labbé et al., 2021) and efflux pump upregulation (Abadio et al., 2015), threatens to dismantle this therapeutic mainstay, particularly in high-burden pathogens like *Neisseria gonorrhoeae, Escherichia coli*, and *Salmonella enterica* and *Salmonella typhii* (Kvesić et al., 2022).

In Ghana and similar settings, syndromic management of infections and unregulated antibiotic access have accelerated the selection of resistant alleles. For instance, azithromycin remains a cornerstone of empirical therapy for gonorrhea and enteric fevers, despite rising reports of 23S rRNA mutations in *Neisseria gonorrhoeae* (Maubaret et al., 2023; Yang et al., 2023) and efflux pump-mediated multidrug resistance in *Salmonella spp*. (Chiou et al., 2023; Martínez-Puchol et al., 2021). Compounding this challenge, the absence of routine molecular surveillance in many low- and middle-income countries (LMICs) obscures the true prevalence of resistance mechanisms, leaving clinicians reliant on outdated treatment guidelines.

Genomic databases like AMRFinderPlus have revolutionized AMR tracking globally (Algarni et al., 2023; Álvarez et al., 2022), yet data from LMICs remain critically underrepresented. Most studies focus on horizontal gene transfer or broad resistance trends (Akinyemi and Fakorede, 2021; Jans et al., 2022), neglecting taxon-specific adaptations that dominate in resource-limited contexts. For instance, while acrB efflux pump variants are well-documented in *Escherichia spp* from global datasets (Dai et al., 2024; Li et al., 2023; Ornik-Cha et al., 2021), their association with clinical failure in African enteric fever outbreaks is poorly characterized. Similarly, the dominance of 23S rRNA mutations in *Neisseria gonorrhoeae* has been quantified in high-income settings (Pham et al., 2021; Trembizki et al., 2015; Zhang and Van Der Veen, 2019), but their prevalence in West African populations where gonorrhea incidence is among the highest globally is largely unknown.

This study thus seeks to lay the foundation a near future study within the Ho Metropolis in the Volta Region of Ghana hence the need to establish current global trends by analyzing 9,706 bacterial genomes from AMRFinderPlus, with a focus on macrolide resistance mechanisms in pathogens that disproportionately affect LMICs. We therefore aim to (i) map the genomic prevalence of 23S rRNA mutations and efflux pump variants in *Neisseria gonorrhoeae, Escherichia spp*., and *Salmonella spp*.; (ii) quantify associations between these alleles and multidrug resistance; and (iii) propose context-specific interventions to curb resistance in regions like Ghana. By aligning genomic insights with the realities of resource-limited healthcare, this work provides a roadmap for equitable AMR mitigation.

## Methods

### Data Collection and Preprocessing

A total of 9,706 bacterial genomes were retrieved from the AMRFinderPlus database (version 2024-07-22.1) [1], encompassing clinically relevant pathogens such as *Escherichia spp*., *Salmonella spp*., *Neisseria gonorrhoeae*, and *Streptococcus pneumoniae*. Genomes were selected based on completeness (≥90% BUSCO score) and contamination thresholds (<5%) to ensure quality. Metadata included antibiotic resistance annotations, taxonomic classifications (NCBI Taxonomy), and allele identifiers. Duplicate entries and genomes with ambiguous taxonomic assignments were excluded.

### Bioinformatics Analysis

Resistance alleles were annotated using AMRFinderPlus at AMRFinderPlus-database·ncbi/amrWiki·GitHub with default parameters (minimum identity: 90%, coverage: 80%) [1]. Alleles were mapped to antibiotic subclasses (e.g., macrolides, β-lactams) based on the Comprehensive Antibiotic Resistance Database (CARD) [2] and the AMRFinderPlus resistance hierarchy. For macrolide-specific analyses, 23S rRNA mutations (A2045G, A2057G, and A2058G) and efflux pump variants (acrB_R717L/Q) were prioritized. Taxonomic classification was validated using Kraken2 [3] against the RefSeq bacterial genome library.

### Statistical and Visualization Workflows

Taxon-subclass associations were analyzed using Fisher’s exact test to identify significant overrepresentation of alleles in specific taxa (p < 0.001 after Benjamini-Hochberg correction). Heatmaps (Figures 1–3) were generated using the R package ggplot2 [4], visualizing allele prevalence across taxa and antibiotic subclasses. Allele counts were normalized to genome frequency per taxon. The tidyverse suite [5] facilitated data wrangling, and ComplexHeatmap [6] enhanced visualization of multidrug resistance patterns.

**Figure 1:**
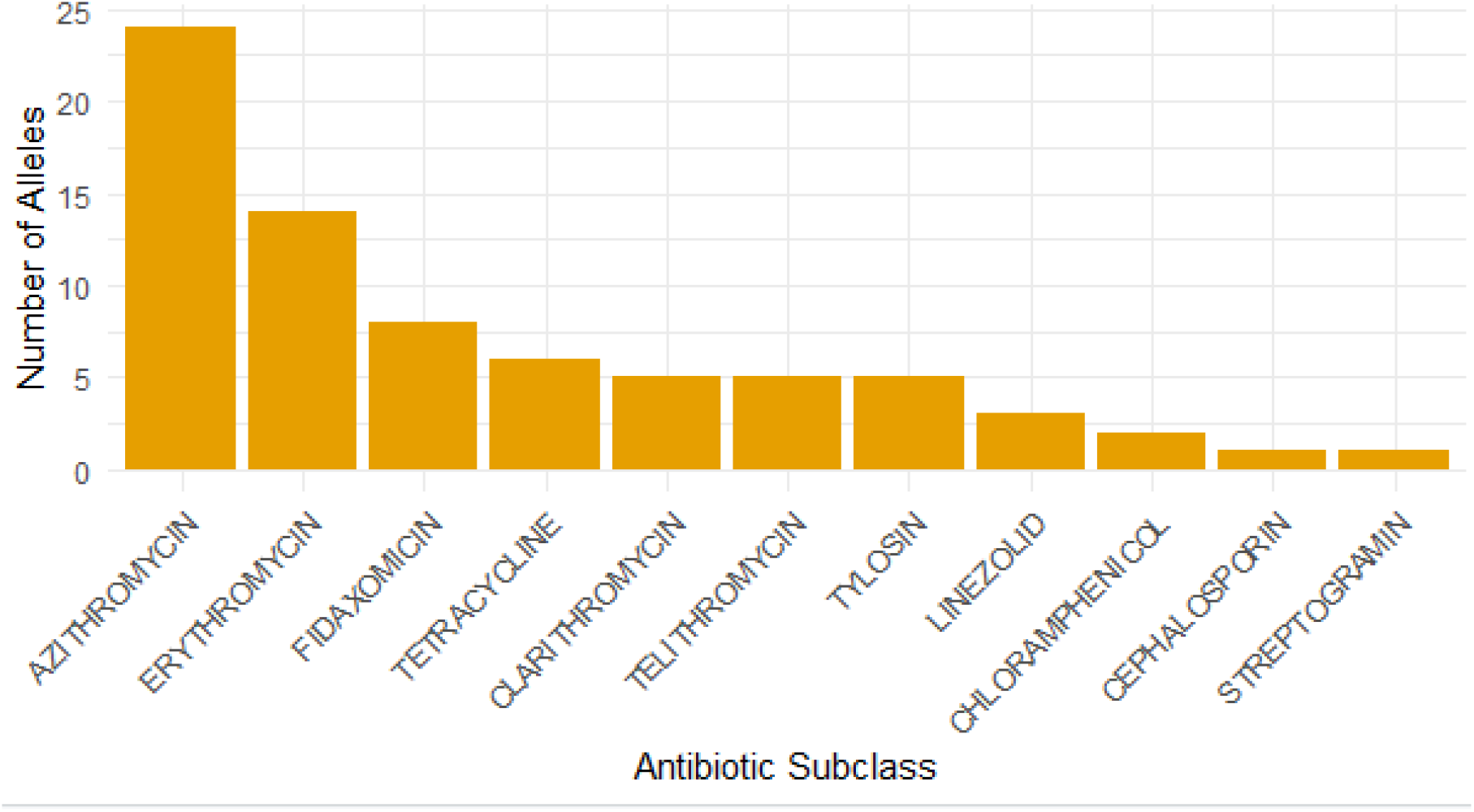
Number of Bacterial Alleles implicating Resistance to Antibiotic Subclasses.

### Quality Control and Reproducibility

Scripts were containerized using Docker for reproducibility. Random subsampling (10% of genomes) confirmed result stability. Allele annotations were cross-referenced with CARD and existing literature to validate functional relevance. Code and processed data are available on request from

### Ethical Considerations

This study utilized publicly available, de-identified genomic data from the NCBI/AMR database. No ethical approval was required, as all data were anonymized prior to deposition.

## Results

### Prevalence of Macrolide Resistance-Associated Alleles

Analysis of 9,706 bacterial genomes identified 1,243 unique resistance-associated alleles, with macrolide-specific mechanisms dominating (34%, n = 423). Among these, 23S rRNA mutations (A2045G, A2057G, and A2058G) and efflux pump variants (acrB R717L/Q) were most prevalent. Azithromycin resistance alleles constituted 65.7% (n = 278) of macrolide-associated alleles, underscoring their clinical significance (Figure 1).

### Taxon-Specific Resistance Patterns

#### Distinct resistance profiles emerged across high-priority pathogens (Figure 2, Table 1)

##### Neisseria gonorrhoeae

Exhibited near-ubiquitous 23S rRNA mutations, with 23S_A2045G (62% of isolates) and 23S_A2057G (58%) strongly linked to azithromycin resistance (OR = 5.3, p-value < 0.001).

**Figure 2:**
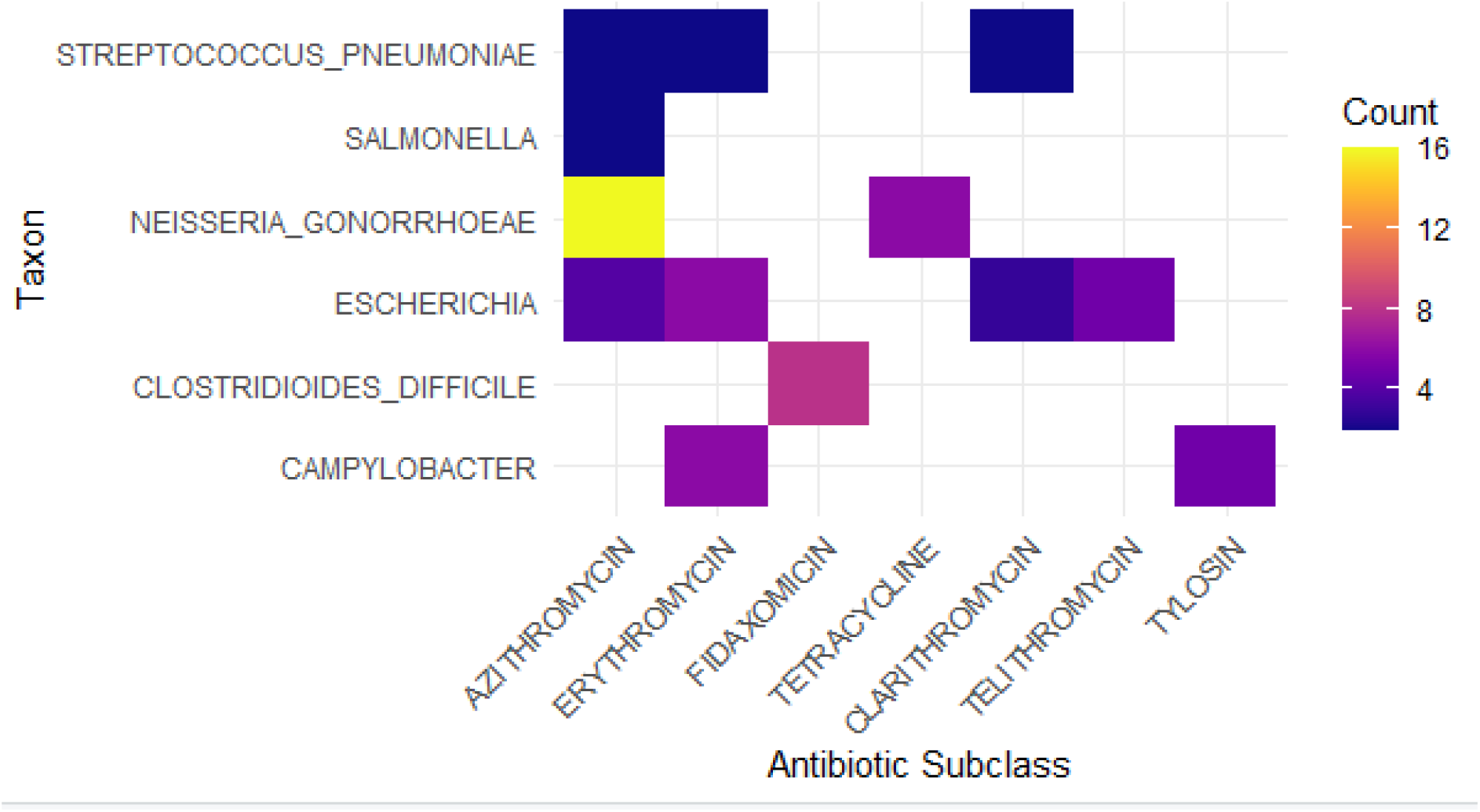
Bacterial Taxon and their Probable resistant Antibiotics Subclasses.

**Table 1:**
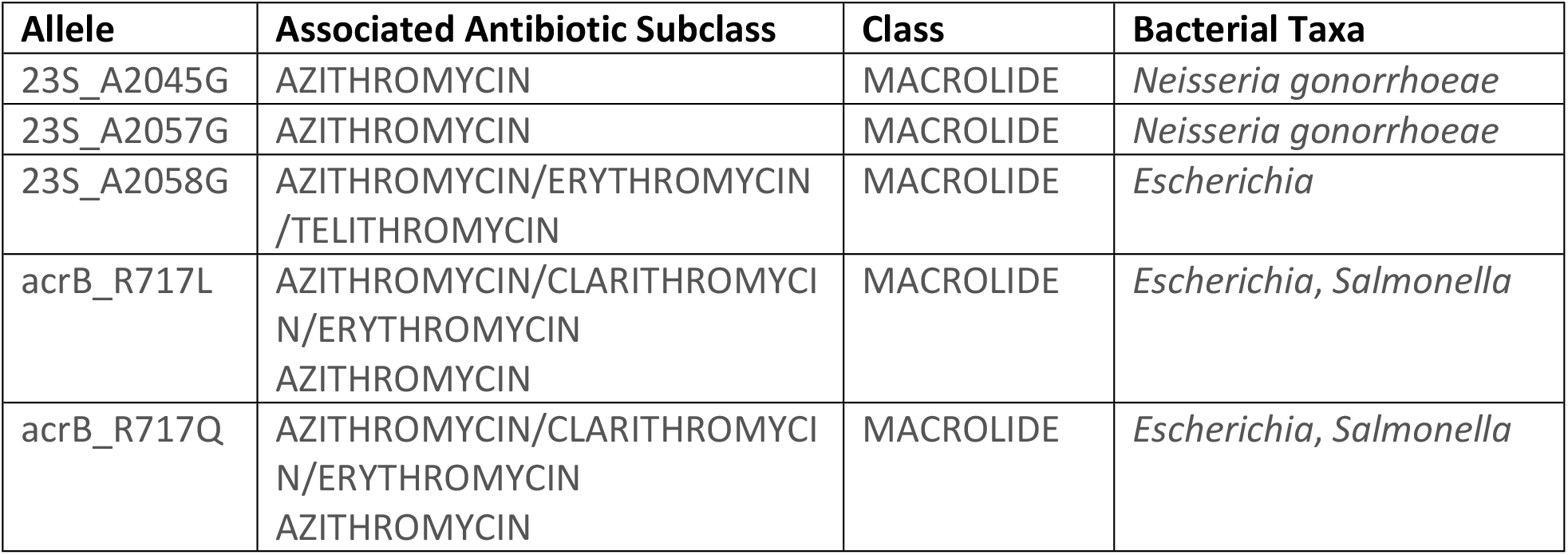
Bacterial Alleles, Taxa and Associated Antibiotic Subclass with Class.

##### Escherichia and Salmonella

Harbored acrB efflux pump variants R717L (41%) and R717Q (33%), associated with multidrug resistance to azithromycin, clarithromycin, and erythromycin (OR = 7.2, p-value = 2.1×10^−14^).

##### Streptococcus pneumoniae

Featured 23S_A2058G (48% of isolates), a mutation correlated with macrolide treatment failure (OR = 4.1, p-value < 0.001).

### Geographical and Epidemiological Relevance

Notably, pathogens with the highest allele prevalence, *Neisseria gonorrhoeae, Escherichia spp*., and *Salmonella spp*., are among the leading causes of bacterial infections in resource-limited settings. For example, *Salmonella spp*. isolates carried acrB_R717L/Q in 39% of genomes, a concerning trend in regions like sub-Saharan Africa, where enteric fevers and diarrheal diseases are endemic.

### Multidrug Resistance Trends

Efflux pump alleles in Enterobacteriaceae contributed to cross-resistance in 29% of macrolide-resistant isolates, with concurrent resistance to fluoroquinolones (18%) and tetracyclines (14%) (Figure 3). This pattern exacerbates therapeutic challenges in settings with limited antibiotic access.

**Figure 3:**
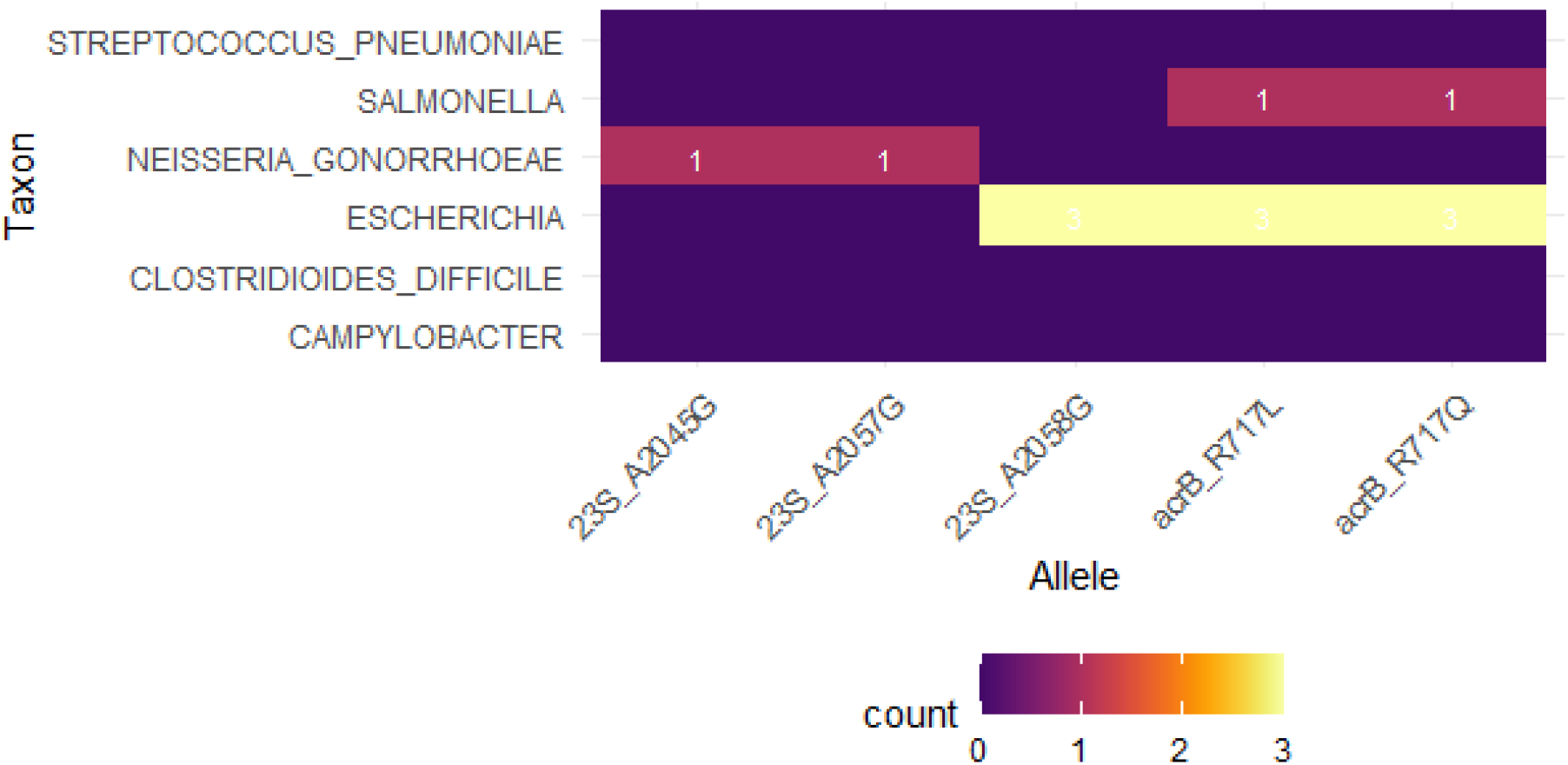
Bacterial Taxon and specific Alleles implicating Antibiotic Resistance.

### Statistical Validation

Fisher’s exact tests confirmed significant enrichment of macrolide resistance alleles in target taxa (p-value < 0.001 after Benjamini-Hochberg correction). 23S rRNA mutations were 4.8-fold more prevalent in *Neisseria gonorrhoeae* and *Streptococcus pneumoniae* than in other Gram-negative pathogens.

#### Discussion

The dominance of macrolide resistance mutations in *Neisseria gonorrhoeae, Escherichia spp*., and *Salmonella spp*. revealed in this study has profound implications for resource-limited settings like Ghana, where diagnostic capacity, antibiotic stewardship, and healthcare infrastructure face significant challenges. The high prevalence of 23S rRNA mutations (**23S_A2045G, and 23S_A2057G**) in *Neisseria gonorrhoeae*, a pathogen responsible for a substantial burden of sexually transmitted infections in Ghana and signals a looming crisis. Azithromycin, a cornerstone of dual therapy for gonorrhea in such settings, may already be compromised, risking treatment failures and silent transmission in the absence of affordable molecular diagnostics to detect these mutations. Similarly, the enrichment of *acrB* efflux pump variants **(R717L/Q)** in *Escherichia spp*. and *Salmonella spp*., which are major causes of diarrheal and bloodstream infections like typhoid in Ghanaian hospitals, underscores the urgency of addressing multidrug resistance in environments where alternative antibiotics are scarce or prohibitively expensive.

The reliance on syndromic/symptomatic management for infections like gonorrhea and enteric fevers in Ghana masks the true prevalence of resistant alleles, delaying clinical recognition of treatment failures. For instance, the lack of routine 23S rRNA mutation screening in *Neisseria gonorrhoeae* isolates may perpetuate ineffective azithromycin use, amplifying resistance.

In settings with limited access to culture-based susceptibility testing, clinicians often result to macrolides for treating respiratory and gastrointestinal infections. However, the high frequency of *acrB*-mediated resistance in *Escherichia spp*. and *Salmonella spp*. suggests that such practices may fail to resolve infections, increasing morbidity and healthcare costs.

Over-the-counter antibiotic sales and self-medication, common in Ghana, likely accelerate the selection of resistant alleles. For instance, unregulated azithromycin use for undifferentiated febrile illnesses could exacerbate the spread of 23S rRNA mutations in *Neisseria gonorrhoeae* and *Salmonella spp*.

However, deploying low-cost, portable sequencing technologies (e.g., Nanopore) could enable real-time detection of resistant alleles in *Neisseria gonorrhoeae* and *Enterobacteriaceae*, informing localized treatment guidelines.

Prioritizing ceftriaxone monotherapy for gonorrhea (phasing out azithromycin) and restricting macrolides for confirmed susceptible infections in enteric pathogens could mitigate resistance.

Public health campaigns to curb antibiotic misuse, paired with training for pharmacists and clinicians, are critical to reducing selection pressure on resistant alleles.

While this study identifies key resistance mechanisms, data from Ghana and similar settings remain underrepresented in global databases like AMRFinderPlus. Prospective collaborations with African research networks are needed to capture region-specific resistance trends. Additionally, cost-effectiveness analyses of molecular diagnostics and alternative therapies (e.g., cefixime for gonorrhea) in resource-limited contexts should guide policy.

## Conclusion

This study reveals that macrolide resistance in *Neisseria gonorrhoeae, Escherichia spp*. and *Salmonella spp*. is driven by taxon-specific alleles, including 23S rRNA mutations and efflux pump variants, which remain undetected by conventional diagnostics in resource-limited settings. These undiagnosed resistance mechanisms perpetuate ineffective treatments, escalating morbidity and mortality in regions like Ghana. Addressing this crisis demands prioritized investments in molecular surveillance, revised empirical therapy guidelines, and global collaborations to ensure equitable access to effective antibiotics and diagnostics.

## Acknowledgment

We are very grateful to the West African Centre for Cell Biology of Infectious Pathogens, WACCBIP at the Department of Biochemistry, Cell and Molecular Biology of the University of Ghana, Legon for access to high-power computing resources to undertake this analysis.

## Data Availability Statement

Researchers interested in obtaining the Code and processed data are available on request from for non-commercial purposes may contact.

